# Exocyst stimulates each step of exocytic SNARE complex assembly and vesicle fusion

**DOI:** 10.1101/2022.01.16.476540

**Authors:** Chanwoo Lee, Dante Lepore, Mary Munson, Tae-Young Yoon

## Abstract

The exocyst is a large multisubunit tethering complex essential for targeting and fusion of secretory vesicles in eukaryotic cells. Although the assembled exocyst complex is proposed to tether vesicles to the plasma membrane and activate the SNARE proteins for membrane fusion, only little is known about the key biochemical steps that exocyst stimulates in the course of SNARE complex assembly, a critical question defining the essential molecular role of the exocyst complex. Here, we use a combination of single molecule and bulk fluorescence assays with purified octameric yeast exocyst complexes to examine the role of exocyst in a reconstituted SNARE assembly and fusion system. We show that the exocyst complex simulates multiple steps spanning from SNARE protein activation to ternary complex assembly, rather than affecting only a specific subset of steps. We also observed that the exocyst has important downstream roles in driving membrane fusion, up to full content mixing of vesicle lumens. Our results suggest that the exocyst complex provides extensive chaperoning functions for the entire process of SNARE complex assembly, presumably using its multi-faceted structure provided by the eight subunits.

## Introduction

The delivery of vesicular cargo between intracellular compartments, and to the outside of a cell, is an essential eukaryotic process underpinning cell growth, motility, signaling, and homeostasis. Many regulatory mechanisms exist to ensure cargo is correctly trafficked to predetermined destinations, preserving cellular organization and compartmentalization of function. Membrane fusion is the last step in any vesicular trafficking pathway and the last chance for regulatory control.

SNARE proteins are the minimal machinery for membrane fusion, and thus essential for cellular trafficking (Weber et al., 1998). SNAREs are transmembrane proteins present on both the vesicle (v-SNAREs) and target membrane (t-SNAREs) and provide the mechanical force for membrane fusion by forming tight four-helix bundle complexes with each other (Antonin et al., 2002; Strop et al., 2008; Sutton et al., 1998; Yoon and Munson, 2018). Each SNARE contributes one or two regions called SNARE motifs to these four-helix bundles; the N- to C-terminal zippering of these SNARE motifs together pulls the vesicle and target membranes into close enough proximity to facilitate membrane fusion (Diao et al., 2012; Gao et al., 2012; Ma et al., 2015; Min et al., 2013; Weber et al., 1998; Zhang, 2017). The four-helix bundle contains a mainly hydrophobic core, except for a polar “zero-layer” in the center, which contains one glutamine (Q) and 3 arginine (R) residues. Each SNARE motif contributes a residue for this zero-layer and are thus named Q- or R-SNAREs (Fasshauer et al., 1998; Yoon and Munson, 2018).

Recombinant SNAREs are capable of inducing liposome fusion *in vitro*, however the observed speed of reconstituted SNARE-mediated fusion is slower than fusion occurs *in vivo* (Li et al., 2016). Additionally, while each SNARE complex appears specific to a particular trafficking step, individual SNARE proteins can interact somewhat promiscuously (Tsui and Banfield, 2000). Therefore, for temporally and spatially accurate cargo delivery, SNARE complex formation and fusion must be tightly controlled by a number of regulatory factors (Brennwald et al., 1994; Fukuda et al., 2000; Gurunathan et al., 2000; Mollard et al., 1997; Nicholson et al., 1998; Schwartz et al., 2017). Furthermore, many of the syntaxin family of t-SNAREs contain auto-inhibitory N-terminal Habc domains that can form intramolecular four-helix bundles with the adjacent SNARE motif (Dulubova et al., 1999). The transition from these auto-inhibitory “closed” conformations to “open” conformations, in which the SNARE motif is released from these intramolecular interactions, is a rate-limiting step for SNARE complex formation (MacDonald et al., 2010; Munson et al., 2000; Nicholson et al., 1998). Many families of SNARE regulatory proteins have previously been identified to function in different trafficking pathways and cell types. Critically, regulators such as the Sec1/Munc18 family proteins and Multi-subunit Tethering Complexes (MTCs) play essential roles, although the mechanistic details are not well understood (Dubuke and Munson, 2016; Ma et al., 2013; Yoon and Munson, 2018).

Exocyst is an essential hetero-octameric MTC that controls exocytosis in eukaryotes including *S. cerevisiae*, and molecular details of its structure have been determined by cryo-EM combined with chemical cross-linking (Ganesan et al., 2020; Heider et al., 2016; Lepore et al., 2018; Mei et al., 2018). Its eight subunits have been shown to directly interact with all three exocytic SNARE proteins (Dubuke et al., 2015; Shen et al., 2013; Sivaram et al., 2005; Yue et al., 2017). The purified N-terminal region of the exocyst subunit Sec3 was shown to bind the exocytic t-SNARE Sso2; this interaction affects the conformation of the Sso2 Habc domain and leads to an increased rate of Sso2 binding to its partner t-SNARE Sec9, as well as stimulates liposome fusion (Yue et al., 2017). Additionally, the purified exocyst subunit Sec6 is able to interact with both the v-SNARE Snc2, and to Sec9 (Dubuke et al., 2015; Shen et al., 2013; Sivaram et al., 2005), although the functional consequences of these interactions are not well understood. In addition, how the full exocyst complex interacts with a plethora of binding partners, and how it regulates SNARE complex assembly and membrane fusion had not been characterized. In particular, identification of the key biochemical step that exocyst accelerates in SNARE-mediated membrane fusion is a crucial question that remains unanswered, which will provide insights into how exocyst regulates this last step of membrane trafficking.

Here, we combined our robust, reconstituted membrane fusion assay containing all three exocytic SNAREs with purified, intact exocyst complexes, and demonstrated that exocyst provides extensive chaperoning functions for the entire course of SNARE complex assembly. Using the single molecule fluorescence resonance energy transfer (FRET) analysis, we observed that exocyst triggers the “open” conformation of Sso1 to increase the rate of initial binary SNARE assembly. We further found that independent to opening of Sso1, exocyst stimulates ternary SNARE complex formation. Finally, we discovered that exocyst can further stimulate membrane fusion downstream of SNARE assembly. Thus, our results suggest that the full exocyst complex may have the potential for stimulating each of the steps of SNARE complex assembly and the downstream membrane fusion.

## Results

### Exocyst facilitates formation of binary Q-SNARE complexes

Sso1 and its paralog Sso2 are yeast exocytic Q-SNARE proteins, which have N-terminal Habc domains that associate with the intramolecular C-terminal SNARE motifs to lock the Sso1/2 protein into a “closed conformation” that prohibits interactions with other SNAREs (Munson, 2015; Munson and Hughson, 2002). We first examined whether the exocyst complex is involved in opening of Sso1, which is a required first step in regulation of exocytic SNARE-mediated membrane fusion (Nicholson et al., 1998). To probe conformational changes of individual Sso1 proteins, we introduced two cysteine residues at a.a. 120 in the Habc domain, and a.a. 150 in the middle of the SNARE motif, respectively. We labeled these cysteines in the purified Sso1 proteins with Cy3 and Cy5 dyes, which make a robust reporter pair for single-molecule fluorescence resonance energy transfer (FRET) (Fig. 1a). The labeling positions were chosen such that when Habc folds back onto the SNARE motif, thereby closing Sso1, the two dyes are brought into molecular proximity to induce high efficiency single-molecule FRET.

**Fig 1.**
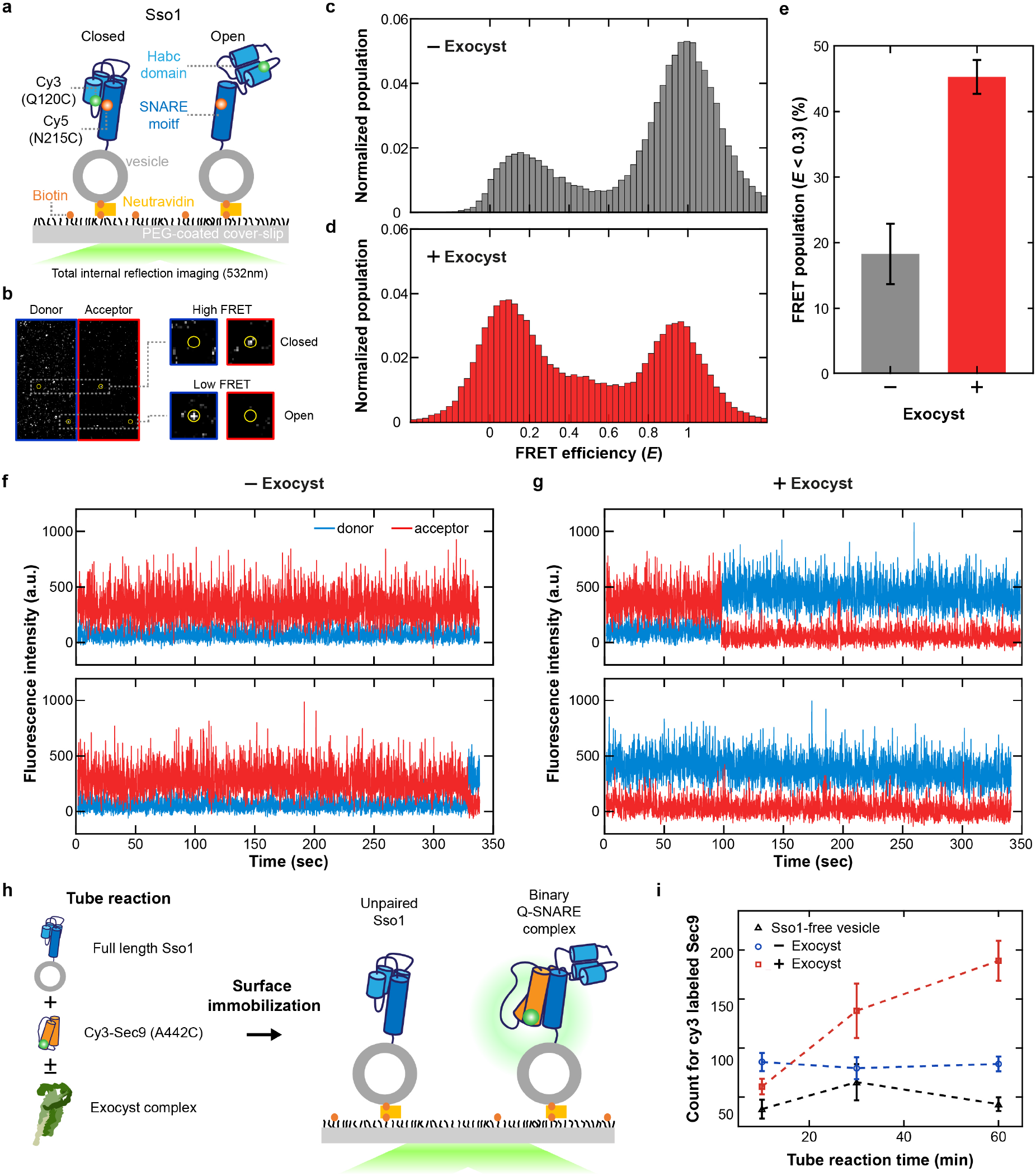
Exocyst enhances formation of the binary Sso1-Sec9 Q-SNARE complex. **a**, Schematics of the single-molecule FRET experiment with Sso1 embedded in vesicles. **b**, Representative images of dual channels under 532nm excitation. Panels with blue and red borders indicate donor (Cy3) and acceptor (Cy5) channels, respectively. Right panels show co-localized areas in each channel marked by yellow circles. The details for confirmation of Cy3 and Cy5 double labeling are in the methods. **c, d**, FRET histogram of double-labeled Sso1 without exocyst (**c**) or with exocyst (**d**). **e**, Low FRET population (*E* < 0.3) of double-labeled Sso1. Error bars show the standard deviation (*n* = 3). **f, g**, Representative traces of donor and acceptor intensity without (**f**) or with (**g**) exocyst. Under 532nm excitation, the acceptor intensity (red) is higher when Cy3 and Cy5 are close and FRET efficiency is high. Donor intensity (blue) is higher when Cy3 and Cy5 are farther away and FRET efficiency is low. Two replicates are shown for each condition. **h**, Schematic diagram of single-molecule binding experiment of Cy3-labeled Sec9 to Sso1-embedded vesicles, with and without exocyst. **i**, Quantification of Cy3-labeled Sec9 bound to Sso1-embedded vesicles, with and without exocyst, over the indicated incubation times. Error bars show the standard deviation (*n* = 5).

To examine the conformation of Sso1 proteins in the absence and presence of exocyst, we reconstituted the FRET pair-labeled Sso1 proteins with their transmembrane domains in vesicles and immobilized these vesicles on an imaging plane of the total internal reflection (TIR) fluorescence microscope (Fig. 1a). We used a low protein-to-lipid ratio of 1:25000 in our vesicle reconstitution, which resulted in most vesicles containing less than one Sso1 protein (Ryu et al., 2015). By measuring the resulting FRET efficiency (*E*), we determined the conformational state (open vs. closed) of Sso1 (Fig. 1b,c). Additionally, we measured photobleaching of both Cy3 and Cy5 dyes to ensure that we only included vesicles containing a single, Cy3 and Cy5 labeled Sso1 protein in our final analysis (Supplementary Fig. 1).

First, we examined the distribution of FRET efficiencies exhibited by single Sso1 proteins alone, and found that most of the Sso1 proteins (i.e., more than 70%) showed the high FRET efficiency values around 1 (Fig. 1c). This observation indicates that the Sso1 proteins are mainly folded in the closed state, an expected observation given the close proximity of the Habc domain and the SNARE motif that are connected by a short linker (Fig. 1a) (Munson et al., 2000). We then determined the FRET efficiency distribution of the same batch of Sso1-reconstituted vesicles after incubation with purified exocyst complex for 15 minutes, a reaction time that allows enough interactions between single Sso1 and exocyst complexes at the exocyst concentration used (10 nM). Remarkably, the population showing the lower FRET efficiency (*E*<0.3) increased at the expense of the high FRET population, reaching 45% of the total populations, compared to ~20% without exocyst (Fig. 1c-e). We interpreted the low FRET efficiency population as Sso1 proteins in an open conformation (Fig. 1a). Thus, this change in the single-molecule FRET signal upon addition of exocyst clearly indicated that exocyst induced a marked shift from closed to open Sso1, with the Habc domain dissociated.

We next examined kinetic transitions recorded in our time-resolved single-molecule FRET traces. For individual Sso1 proteins without exocyst, most of the FRET traces were largely stationary, remaining in the high FRET values with very seldom transitions to lower FRET values (Fig. 1f). We found that the addition of exocyst resulted in many of the traces residing in the low FRET states (thus, the open states) when we started the recording. However, the presence of exocyst did not significantly increase the frequency of transitions among the open and closed states, with only a small proportion of the FRET traces showing a transition to the other FRET state (Fig. 1g, upper). Although we did not carry out detailed kinetics analysis because of the scarcity of the transition events within our observation time of ~300 s, these observations indicated that in the absence of exocyst or other factors, the Habc domain mostly remains bound to the SNARE motif, thereby closing Sso1, rather than frequently opening and closing. Our data also suggested that after exocyst induces the open conformation Sso1, exocyst remains bound to Sso1 to maintain Sso1’s opened state.

Finally, we examined whether opening of Sso1 would facilitate binding to its partner Q-SNARE Sec9 (our construct contains a.a. 416–651, which are the essential residues of Sec9 *in vivo*) (Brennwald et al., 1994). We mixed Sso1-reconstituted vesicles (wild-type (WT), unlabeled Sso1) with Cy3-labeled Sec9 protein (Fig. 1h, left). After incubating the binding reaction in bulk, we quenched the reaction via dilution and immobilized the Sso1-vesicles on the surface (Fig. 1h, right). By counting single-molecule Cy3 spots, we quantified the amount of Sec9 pulled down with Sso1-vesicles over time (Fig. 1h,i). We observed an increase in the number of Cy3 spots with reaction time, exclusively in the presence of exocyst during the bulk reaction (Fig. 1i). Without either exocyst or Sso1 in the vesicle membranes, the Cy3 spot numbers remained at the background level, an indication of specificity of the reconstituted reaction. These observations demonstrated that opening of Sso1 by exocyst leads to enhanced formation of binary Q-SNARE complexes composed of Sso1 and Sec9.

### Exocyst facilitates formation of ternary SNARE complexes

Formation of the ternary SNARE complex requires binding of the R-SNARE Snc1/2 to the binary Sso1:Sec9 Q-SNARE complex. Because we observed that exocyst effectively induced Sso1 opening and formation of binary Q-SNARE complexes, we asked whether exocyst is also involved in the later stage of ternary SNARE complex formation. Unlike the binary Q-SNARE complexes that form via binding of soluble Sec9 to membrane-bound Sso1, the association between R- and Q-SNAREs occurs while both of them are anchored in the opposite membranes, forming so-called *trans*-SNARE complexes. Previous results based on MD simulation suggested that apposition of two membranes into such a nanometer proximity brings about substantial repulsion force amounting to tens of pN (Bykhovskaia et al., 2013; Shon et al., 2018). We thus examined whether exocyst could facilitate *trans-*SNARE complex formation with both the R- and Q-SNAREs reconstituted in separate membranes.

To this end, we prepared Sso1-reconstituted vesicles with one modification—we truncated the inhibitory Habc domain from Sso1 such that we could determine the kinetics of binding among R- and Q-SNAREs without complications from the preceding step of Sso1 opening. After reconstitution of Habc-truncated Sso1 in vesicle membranes, we added an excess of Sec9 protein (at 0.5 μM) to induce formation of the binary Q-SNARE complexes. We prepared another group of vesicles reconstituted with R-SNAREs (Snc2), and introduced these R-SNARE vesicles into the reaction tube. Finally, we selectively immobilized the Q-SNARE vesicles on the imaging plane of the single-molecule fluorescence microscope because only the Q-SNARE vesicle membrane contained lipids with biotinylated headgroups.

We labeled the R-SNARE vesicles with the lipophilic dye DiD such that the formation of SNARE complexes between R- and Q-SNARE vesicle membranes appeared as stable, diffraction-limited fluorescence spots in the DiD channel (Fig. 2a, note that the degree of membrane fusion between R- and Q-SNARE vesicle membranes was not assessed in this experiment). We first examined SNARE complex formation in the absence of exocyst and observed an increase in the DiD spot numbers with reaction time (Fig. 2b). When we omitted Sec9, the DiD spot number remained at the background level through the reaction time we studied, indicating that the observed increase was specific for SNARE complex formation (Fig. 2b). Remarkably, when adding 10 nM exocyst in our vesicle-vesicle reaction, we observed an increase in DiD spot numbers compared to those observed without exocyst over the entire reaction time, suggesting that exocyst significantly facilitated assembly of SNAREpins and concomitant vesicle-vesicle complexes on our single-molecule fluorescence microscope (Fig. 2b). We repeated the same reaction with full-length Sso1, including the inhibitory Habc domain, and found a similar increase in the DiD spot numbers with exocyst (Fig. 2c). This observation indicates that independent of the role of Sso1 opening, exocyst can stimulate ternary Sso1:Sec9:Snc2 SNARE complex assembly from pre-formed binary Sso1:Sec9 complexes.

**Fig 2.**
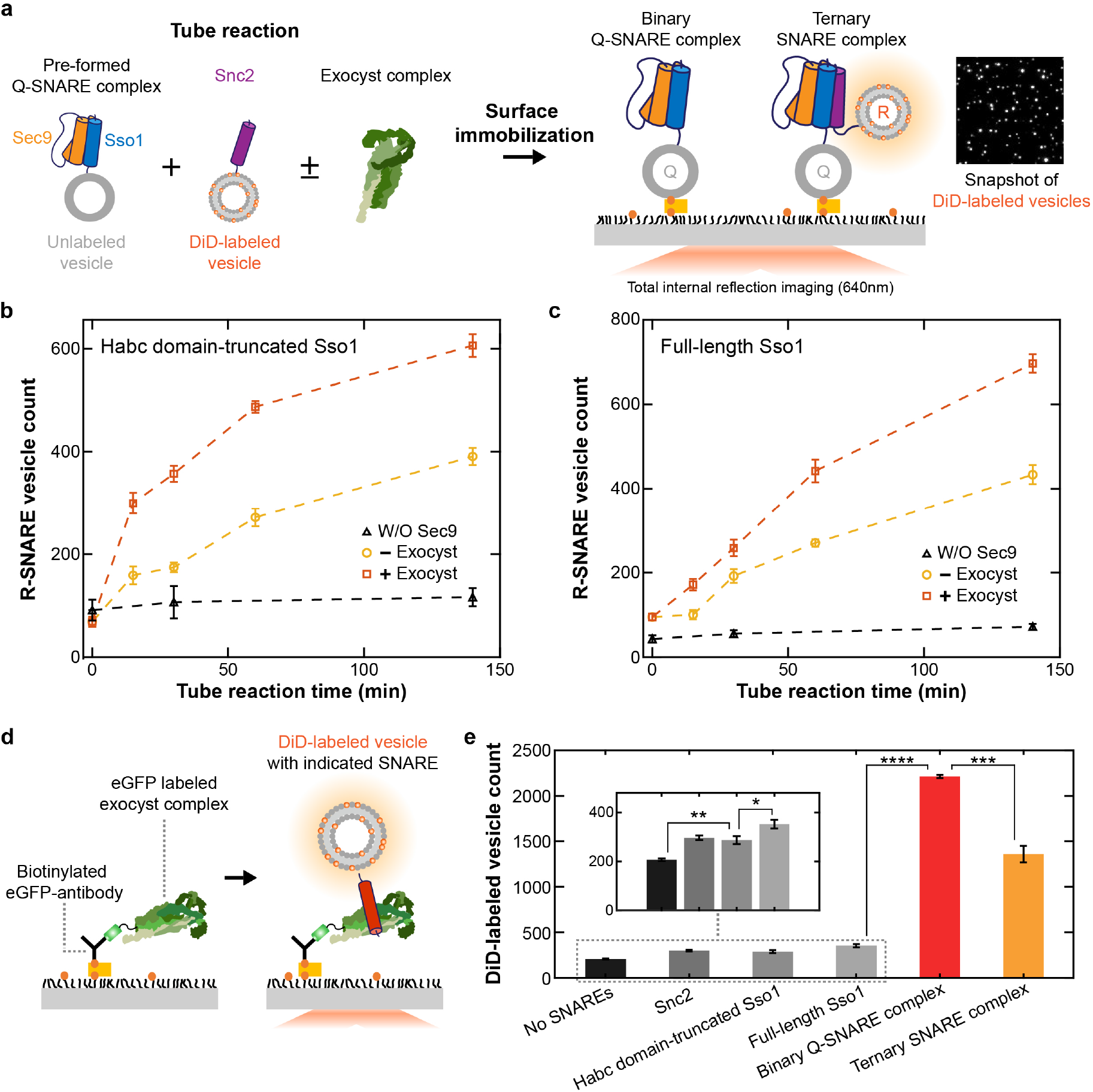
Exocyst stimulates ternary SNARE complex assembly independent of Sso1 opening. **a**, Schematic diagram shows the preparation of mixtures and immobilized vesicles. Pre-formed Q-SNARE complexes were obtained by mixing Sso1 with Sec9 for 1hr at 30 °C before incubation with exocyst and DiD-labeled R-SNARE vesicles in the tube reaction. Biotinylated binary Q-SNARE vesicles were immobilized on the surface. Right panel shows a snapshot of DiD-labeled R-SNARE vesicles. The fluorescence signal (white spot) is generated if the R-SNARE vesicle is immobilized by SNARE complex formation. **b, c**, The number of R-SNARE vesicle over incubation time after being pull-down of Q-SNARE vesicle onto the PEG-coated surface. Habc domain-truncated Sso1 was used for (**b)** and full-length Sso1 was used for (**c)**. Error bars show the standard deviation (*n* = 3). **d**, Schematic diagram of experiments for measuring interactions between exocyst and SNARE proteins. Exocyst complexes and DiD-labeled vesicles were injected into the chamber **e**, Quantification of the number of DiD-labeled vesicles containing SNARE constructs as indicated, which interact with immobilized exocyst complexes. Asterisks represent the level of significance based on two sample t-test. Based on P-value (P), asterisks mean the following. * for P≤0.05, ** for P≤0.01, *** for P≤0.001 and **** for P≤0.0001. Inset shows the magnified view of first four SNARE constructs. Error bars represent the standard deviation (*n* = 5).

Collectively, our results indicate that the exocyst complex can interact with SNARE proteins in almost every assembly state, ranging from single proteins to full ternary SNARE complexes. For example, our single-molecule FRET data suggest that exocyst interacts with individual Sso1 proteins to induce an Habc-domain opened state (Fig. 1). Our single-vesicle immobilization assay suggests that exocyst also interacts with the binary Q-SNARE complexes to facilitate formation of ternary SNARE complexes (Fig. 2a-c). To test our hypothesis in a more quantitative fashion, we constructed an assay that directly determines the interactions between exocyst and SNAREs in various assembly states. For these experiments, we purified exocyst complexes tagged with enhanced green fluorescent protein (eGFP; eGFP fused to the N-terminal end of Exo84) and immobilized these tagged exocyst complexes on the surface using anti-eGFP antibodies (Fig. 2d). We reconstituted SNARE proteins—in different assembly states—into vesicles and reacted these proteo-liposomes (at 40 μM, in terms of lipid concentration) with surface-immobilized exocyst complexes for 10 minutes. Because the vesicle membranes were labeled with DiD, an interaction between SNAREs and exocyst complexes on the surface led to an appearance of single vesicle spots as in Fig. 2a.

When individual Q- and R-SNAREs were reconstituted in vesicle membranes, we observed rather weak but significant interactions with exocyst (Fig. 2e). The interaction became slightly stronger when Sso1 retained the Habc domain, indicating that the Habc domain itself is a part of the binding interface for exocyst and thereby strengthens the interaction (Fig. 2e, inset). Remarkably, when we assembled binary Q-SNARE complexes on vesicle membranes (composed of Habc-truncated Sso1: Sec9), the interaction was increased almost by an order of magnitude (Fig. 2e). This data is consistent with the observed strong stimulatory effect of exocyst on ternary SNARE complex formation (Fig. 2b). When we reconstituted the full ternary SNARE complexes, the exocyst also showed a substantial level of interaction (Fig. 2e), indicating that exocyst would remain bound to the SNARE complexes and continue to exert its effects during the last stages of membrane fusion.

### Exocyst boosts SNARE-mediated membrane fusion

We next examined whether exocyst’s stimulation of protein-protein interactions indeed leads to functional stimulation that enhances SNARE-mediated membrane fusion. We first re-confirmed that two types of vesicles, reconstituted individually with either Snc2 or with Habc-truncated Sso1 plus Sec9, showed robust lipid mixing as we previously determined (Fig. 3a,b) (Yoon et al., 2006). The extent of lipid mixing was measured by labeling each group of vesicles with either DiI or DiD and monitoring the increase in FRET between these lipophilic dyes. When full-length Sso1 protein was used instead in our lipid mixing assay, we observed almost no membrane fusion (Fig. 3c). We attributed this lack of lipid mixing activity to the closed state of Sso1; lipid mixing activity could be rescued by introducing point mutations to Sso1 that keep Sso1 in a constitutively open state (Supplementary Fig. 2a) (Munson and Hughson, 2002).

**Fig 3.**
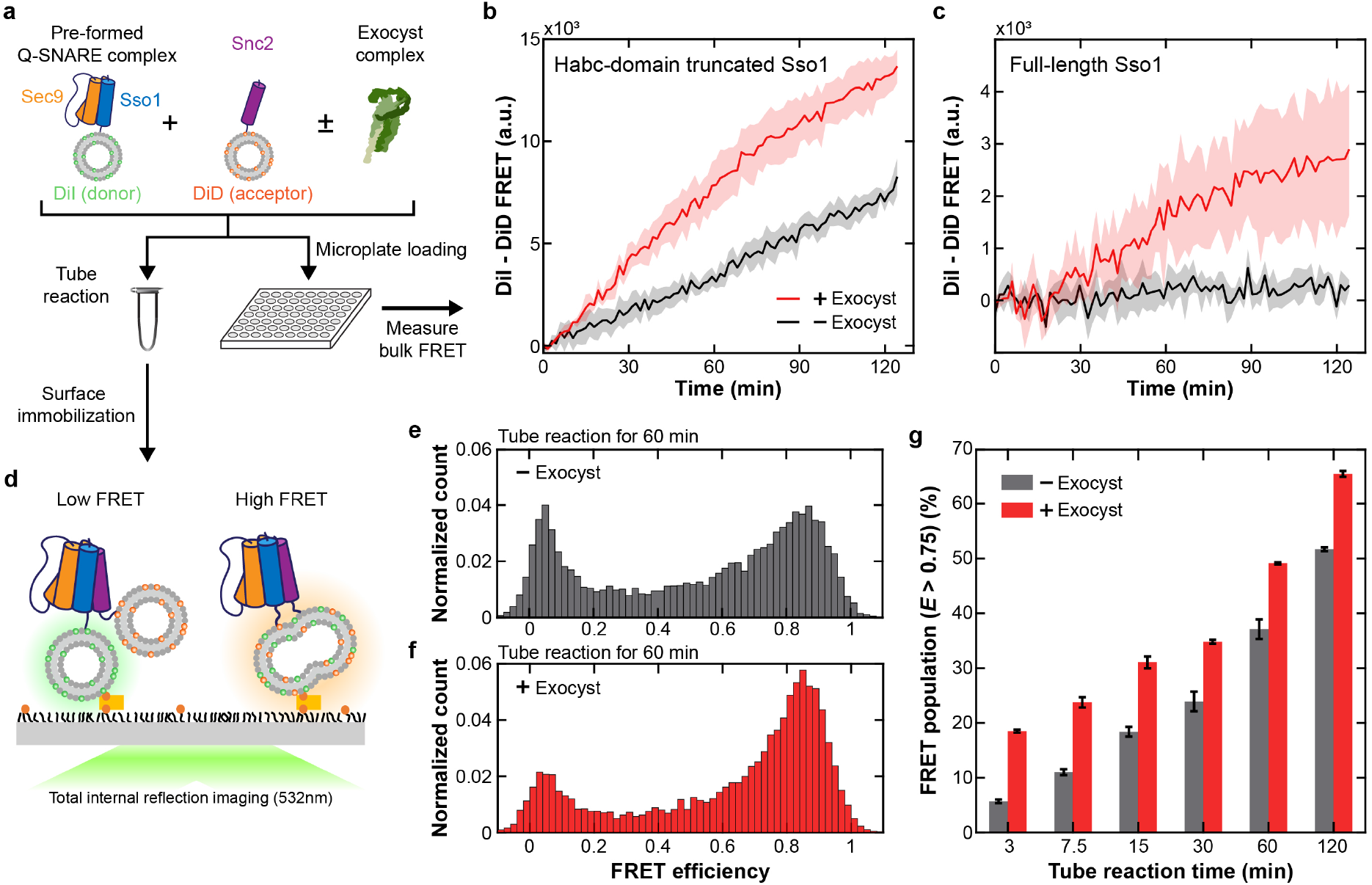
Exocyst accelerates SNARE-mediated membrane fusion. **a**, Schematic diagram of mixture used for lipid mixing experiment. The binary Q-SNARE complex is pre-formed by incubating vesicles containing Sso1 with Sec9. Then, vesicles with Snc2 and exocyst were both added to the pre-mixture. **b**, **c**, DiI-DiD bulk FRET signals generated by lipid mixing of Q-SNARE and R-SNARE vesicle. Habc domain-truncated Sso1 was used for (**b)** and full-length Sso1 was used for (**c)**. The lines represent the mean and shaded area shows the standard deviation from three independent experiments. **d**, Schematics of two cases illustrating low and high FRET signals. **e**, **f**, FRET histogram of single-molecule vesicle fusion experiment without (**e**) or with (**f**) exocyst after incubation for 60 minutes. **g**, Ratio of vesicles with high FRET (0.75 < *E*) to the total vesicles immobilized on the surface over time. Error bars show the standard deviation (*n* = 3).

To examine whether exocyst stimulates this lipid mixing activity, we added exocyst to both types of fusion reactions that were mediated by Habc-truncated or full-length SNARE complexes, respectively. Remarkably, we found that the presence of exocyst markedly enhanced both reactions (Fig. 3b,c). In particular, the increase was more substantial for the fusion reaction driven by the full-length SNARE complexes, as the amount of lipid mixing was minimal in this case without exocyst (Fig. 3c). We further checked whether the enhanced FRET signal between DiI and DiD resulted from genuine lipid mixing between single vesicles. To this end, we immobilized fusion products of the reaction shown in Fig. 3b,c on the imaging plane of our single-molecule fluorescence microscope and examined the fluorescence intensity of single vesicle-vesicle complexes (Fig. 3d). Many of the fusion products showed FRET values centered around 0.8, a high FRET efficiency expected for fully merged vesicles (Fig. 3e,f and Supplementary Fig. 2b,c). Importantly, we found that the proportion of these high FRET populations were consistently larger in the presence of exocyst for all reaction times (Fig 3g).

Finally, because several previous reports indicated that lipid mixing activities are not always translated into corresponding activities at the content mixing level (Chanturiya et al., 1997; Jun and Wickner, 2007), we wondered whether the exocyst stimulatory effect could be observed in an assay measuring the mixing of contents between single vesicles. To this end, we took advantage of 5(6)-carboxyfluorescein (CF) dequenching to monitor fusion between two single vesicles, which would double the vesicle lumen volume and increase the fluorescence signal in a high mM range (Fig. 4a and Supplementary Fig. 3). We encapsulated 100 mM of CF inside Snc2-reconstituted vesicles and induced fusion reactions of these vesicles with unlabeled Sso1 vesicles (Habc-truncated) and Sec9, for 10 minutes. Similarly to experiments in Fig. 3d, we then immobilized the fusion products on the single-molecule fluorescence microscope and observed CF fluorescence intensities exhibited by individual reactions (Fig. 4a). We found the CF fluorescence intensity distribution showed a pronounced peak in the low intensity range followed by a long tail in higher intensity regions (Fig. 4b). When compared with negative-control samples produced via direct immobilization of the CF-quenched Snc2-reconstituted vesicles, the products of the fusion reaction on surface showed increased populations belonging the high-fluorescence tail, accompanied by a decrease in the peak (Fig. 4b). We next repeated this fusion reaction in the presence of exocyst (Fig. 4c). Of note, a substantially higher portion of fusion products appeared in the high-fluorescence tail when compared to the reaction without exocyst (Fig. 4c,d). These observations indicate that exocyst also stimulated mixing between luminal contents, resulting in complete merging of the vesicles.

**Fig 4.**
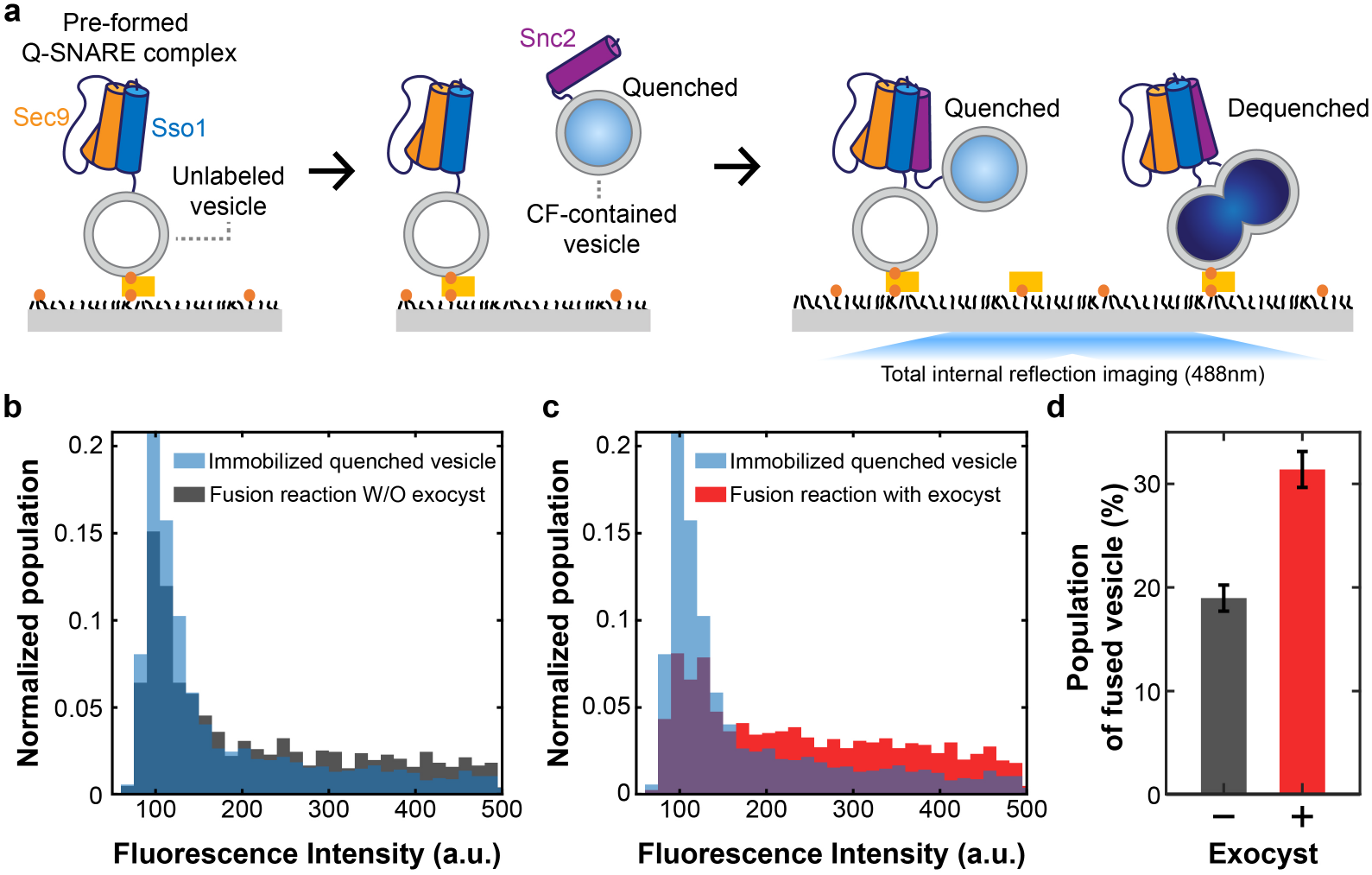
Effect of exocyst on full fusion. **a**, Schematic diagram of the content mixing experiment using 5(6)-carboxyfluorescein (CF) dequenching. Fluorescence emission of self-quenched CF is enhanced when its concentration decreases due to full fusion with an unlabeled vesicle. The Habc domain-truncated Sso1:Sec9 binary Q-SNARE complexes were embedded in unlabeled vesicles and the R-SNARE Snc2 was embedded in the quenched vesicles. **b**, **c**, Intensity histograms of quenched vesicles after incubation for 10 minutes without (**b**) or with (**c**) exocyst in the presence of bound unlabeled vesicles. The data labeled ‘quenched vesicle only’ means that the data was obtained without mixing with unlabeled vesicles. **d**, Ratio of dequenched vesicles due to full fusion to the total CF-containing vesicles immobilized on the surface. Error bars show the standard deviation (*n* = 3).

## Discussion

Many of the regulators of exocytosis have been identified through genetic and biochemical analyses in budding yeast, but details of their functions are not well understood. In particular, exocyst is a huge, hetero-octameric tethering complex of the CATCHR family. Although a moderate resolution structure of the intact yeast exocyst was recently solved (Mei et al., 2018), exocyst mechanisms of action remain to be determined. By building multiple assays for careful biochemical and biophysical dissection of the pathways involved, we showed that addition of purified yeast exocyst complexes to reconstituted exocytic SNARE-mediated vesicle fusion assay stimulates both SNARE complex assembly and membrane fusion.

### Exocyst enhances formation of the binary Sso1-Sec9 Q-SNARE complex

Purified endogenous exocyst complexes from S. cerevisiae were tested in our reconstituted minimal SNARE assembly assays. We showed that exocyst functions to stimulate release of the Habc inhibitory domain of Sso1 to form an open conformation, which facilitates the rate of binary Sso1-Sec9 Q-SNARE complexes. As had previously been observed for a small N-terminal domain of the exocyst subunit Sec3 and Sso2 (Yue et al., 2017), exocyst interacts directly with Sso1 and catalyzes formation of an open Sso1 conformation (Fig. 1). Our single molecule FRET assay captured the numbers of individual open vs. closed Sso1 molecules, and showed that the number of open Sso1 molecules was greatly enhanced by exocyst. Our data also suggests that the exocyst complex remained bound to open Sso1 and can prevent its re-closing. This exocyst:open Sso1 complex, in turn, led to increased binding of Sso1 to Sec9 (Fig. 1h, i). In contrast to Sec3N, which did not appear to remain bound to binary Sso2:Sec9 complexes (Yue et al), exocyst binds stably to both binary Sso1:Sec9 and ternary Sso1:Sec9:Snc2 SNARE complexes (Fig 2e). In addition to Sso1 binding, exocyst can also interact with individual Sec9 and Snc2 SNAREs, albeit with weaker affinities than it has for SNARE complexes (Fig. 2e). These data are consistent with previous binding studies using purified subunits of exocyst: Sec3N-Sso2 (Yue et al., 2017), and binding of Sec6 to both Sec9 and Snc2, as well as to assembled binary and ternary SNARE complexes (Dubuke et al., 2015; Shen et al., 2013; Sivaram et al., 2005). Curiously, previous data using purified Sec6 and cytosolic domains of SNAREs (Dubuke et al., 2015) suggested that binding of Sec6 to ternary SNARE complexes was tighter than to binary; our opposite result here may more closely represent the native binding interaction, with fully assembled exocyst complexes and membrane-anchored SNAREs.

### Exocyst accelerates SNARE-mediated membrane fusion

Opening Sso1 to increase the rate of binary SNARE complex assembly had several expected consequences—the rate of ternary SNARE complex formation was increased (Fig. 2 b,c) and the rate of membrane fusion was accelerated (Fig. 3). Surprisingly, however, exocyst showed an increased rate of ternary SNARE complex assembly even when the Habc domain of Sso1 was not present (Fig. 2b). Given the time of pre-incubation of Habc-truncated Sso1 with Sec9, prior to addition of Snc2 R-SNARE vesicles, we expect that binary SNARE complex formation would be complete (Nicholson et al., 1998); therefore, the exocyst appears to play a role downstream of binary complex formation, possibly recruiting or tethering vesicles together *in vitro*, analogous to tethering vesicles at the plasma membrane. In addition, or alternatively, exocyst may align or “template” binary complexes together with Snc2 to increase their rate of interaction, similar to the function of the lysosomal/vacuolar Vps33 protein, a Sec1/Munc18 family member and component of the HOPS multisubunit tethering complex, in positioning its cognate SNAREs to accelerate assembly and fusion (Baker et al., 2015; Song et al., 2020).

Following binary Q-SNARE assembly in the presence of full-length Sso1 (with Habc), the rate of membrane fusion was nominal. Upon addition of exocyst, this rate significantly increased. An analogous accelerated fusion rate was also observed with Habc-truncated Sso1, although the difference +/− exocyst was not as dramatic, presumably because binary complexes quickly assemble in the absence of the inhibitory Habc domain (Fig. 3). In both experiments, exocyst stimulates fusion at a step after binary Q-SNARE complex assembly. Our controls and contents-mixing assay indicate that the observed acceleration of membrane fusion by exocyst represents full fusion of vesicles mediated by the exocytic SNAREs (Fig. 4). These findings are supported by data indicating that exocyst tethering of vesicles to the plasma membrane can affect the mode of fusion in mammalian cells (An et al., 2021).

### Conclusions

Our data revealed several functions for exocyst complexes in SNARE-mediated membrane fusion: a) stimulation of binary SNARE complex assembly by releasing the inhibition of Sso1’s N-terminal Habc domain; b) stimulation of ternary SNARE complex assembly, through an as yet undetermined mechanism; and a possible third function in c) stimulation of fusion through exocyst interaction with ternary SNARE complexes. This last putative function, which we are currently unable to dissect independently from ternary SNARE complex assembly in our assays, has been proposed for several SNARE binding proteins, such that binding of a large partner to SNARE complexes could mechanically affect membrane curvature to stimulate fusion (D’Agostino et al., 2017). Binding of a partner to ternary SNARE complexes could explain the stimulation of SNARE-mediated fusion observed by addition of the yeast Sec1 protein (Scott et al., 2004), which also binds ternary SNARE complexes (Carr et al., 1999; Togneri et al., 2006). Curiously, Sec1 also binds exocyst (Morgera et al., 2012; Wiederkehr et al., 2004), although it is currently unclear if Sec1 and exocyst function together or separately to regulate fusion in vivo. A complete understanding of the yeast exocytic tethering and fusion mechanism requires future reconstitution with many additional factors for recognition, tethering and exocyst activation, including the Rab GTPase Sec4 (Guo et al., 1999), Rho GTPases such as Cdc42 and Rho3 (Adamo et al., 1999), the regulator Sro7/Sro77 (Lehman et al., 1999) and others. Moreover, mechanistic details will be elucidated through further high resolution structural and biophysical studies of the different binding interactions and potential conformational changes.

## Methods

### Protein constructs

Full length Sso1 WT (a.a. 1-290), Habc domain-truncated Sso1(a.a. 185-290) and Snc2 WT (1-115) were cloned into pGEX with an N-terminal GST-tag and a thrombin cleavage site. Sec9 was cloned into pET28a with a C-terminal His-tag. For single-molecule fluorescence assay, mutated versions of Sso1 (Q120C/N215C) and Sec9 (A442C) were used. For the open form of Sso1, the Sso1-Open1 mutations V84E/K95E/Y148A were introduced into the full-length construct (Munson and Hughson, 2002)

### Expression and purification of SNARE proteins

Sso1 and Snc2 were expressed in Rosetta (DE3) pLysS competent cell (Novagen). The cells were grown in LB medium at 37 °C until the OD_600_ reached ~0.7. Then, IPTG (0.5mM) was added to induce protein expression. After 16 h at 18 °C, the cells were collected by centrifugation and resuspended in lysis buffer (40mM Tris pH 8.0, 150mM NaCl, 0.5% Tx-100, 0.4% sarcosyl, 1x protease inhibitor cocktail (genDEPOT)). The cells were lysed by sonication and nutated for 1 h to solubilize the protein. After centrifugation at 17,000×g for 30 min, the supernatant was nutated with glutathione sepharose 4b resin (GE healthcare) for 4 h. The resin was washed with wash buffer (40mM Tris pH 8.0, 150mM NaCl, 0.2% Tx-100) and detergent-free buffer (25mM HEPES pH 7.4, 100mM KCl). Elution was done using elution buffer (25mM pH 7.4 HEPES, 100mM KCl, 1 % N-octyl-β-D-glucoside (OG)) with thrombin for cleavage of the GST-tag.

The same process was used for expression of Sec9. The cells were collected by centrifugation and resuspended in lysis buffer (25mM Tris pH 8.0, 350mM NaCl, 0.5% Tx-100, 20mM imidazole, 1x protease inhibitor cocktail). The cells were lysed by sonication. After centrifugation at 17,000×g for 30 min, the supernatant was nutated with Ni-NTA agarose resin (QIAGEN) for 1 h. The resin was washed with wash buffer (25mM Tris pH 8.0, 300mM NaCl, 0.1% Tx-100, 20mM imidazole) and detergent free buffer (25mM Tris pH 7.4, 300mM NaCl, 20mM imidazole). Elution was done using high imidazole buffer (50mM HEPES pH 7.4, 150mM NaCl, 500mM imidazole). For size exclusion chromatography, the elute was loaded onto Superdex 200 16/600 column (GE healthcare) equilibrated with 50mM HEPES pH 7.4, 150mM NaCl, 5% glycerol. All purification processes were done at 4 °C. The concentration of proteins was determined by Bradford assay.

### Purification of endogenous yeast exocyst complexes

Exocyst complexes were purified using a genomically incorporated Protein-A tag on the C-terminus of the exocyst subunit Sec15 (MMY1075). For experiments with eGFP labeled exocyst, an eGFP was incorporated on the N-terminus of Exo84 (MMY1643). Yeast cells expressing endogenous exocyst subunits (both with and without protein tags) were grown in YPD media to an OD_600_ of 1.0-1.5. Cells were harvested by centrifugation at 3,000 x g for 5 minutes and cell pellets washed with sterile water. After washing, and a final centrifugation spin for 15 minutes at 3000 x g to remove excess liquid, cell pellets were pushed through a syringe into liquid nitrogen to create frozen yeast noodles. Noodles were cryo-milled with a planetary ball mill grinder at liquid nitrogen temperatures as previously described (Heider et al., 2016).

Cryo-lysed cells were resuspended in lysis buffer (40mM Tris pH 8.0, 200mM sodium citrate) with 1× complete Mini EDTA-free protease-inhibitor solution (Roche Life Science). After centrifugation at 14,000×g for 10 min at 4 °C, rabbit IgG-conjugated magnetic beads (Dynabeads M-270 epoxy (Invitrogen), rabbit IgG (Merck), (Domanski et al., 2012)) were added to the supernatant and the mixture was incubated for 1 h at 4 °C. The beads were collected on a magnetic rack and washed with lysis buffer. The beads were then resuspended in lysis buffer with Pre-Scission protease and incubated for 1.5 h at 4 °C for elution (Supplementary Fig. 5).

### Reconstitution of vesicle with purified SNAREs

All lipids were purchased from Avanti Polar Lipids. 1-palmitoyl-2-oleoyl-glycero-3-phosphocholine (POPC), 1,2-dioleoyl-sn-glycero-3-phospho-L-serine (DOPS), 1,2-dioleoyl-sn-glycero-3-phosphoethanoleamine (DOPE), 1,2-dioleoyl-sn-glycero-3-phosphoethanolamine-N-(cap biotinyl) (Biotinyl CapDPPE) and ergosterol (Sigma Aldrich) were used and ‘%’ in this method section means molar ratio of lipid. For fluorescent labelling, 1,1’-Dioctadecyl-3,3,3’,3’-Tetramethylindocarbocyanine Perchlorate (DiI) and 1,1’-Dioctadecyl-3,3,3’,3’-Tetramethylindodicarbocyanine (DiD) were used as fluorescent lipophilic tracers (Invitrogen). Lipids and reagents were mixed and dried to make the lipid film by desiccating with vacuum pump for at least 3 h. For single-molecule fluorescence assays and single vesicle fusion assays, a lipid composition of 84.3% POPC, 15% DOPS, 0.7% Biotinyl CapDPPE was used for Q-SNARE vesicle and 84.8% POPC, 15% DOPS, 0.2% DiD for R-SNARE vesicle. For vesicle fusion and content mixing assays, Q-SNARE vesicles and R-SNARE vesicles had different lipid compositions: 44.3% POPC, 15% DOPS, 20% DOPE, 18% ergosterol, 0.7% Biotinyl CapDPPE, 2% DiI for Q-SNARE vesicles and 55% POPC, 15% DOPS, 20% DOPE, 8% ergosterol, 2% DiD for R-SNARE vesicles. POPC replaced DiI when fluorescent labelling was not necessary. The mixed lipid film was rehydrated in fusion buffer (25mM HEPES pH 7.4, 100mM KCl) with 3% OG to make a solution with a lipid concentration of 15mM. This solution and purified SNARE (Sso1 or Snc2) were mixed at the desired protein to lipid ratio (1:25000 for the single-molecule FRET experiments,1:1000 for the single DiD-labeled vesicle experiment and 1:250 for fusion assay). For the reconstitution of proteo-liposomes, the protein and lipid mixtures were diluted three-fold with fusion buffer to lower the concentration of OG below the critical micelle concentration. Also, the concentration of lipid is determined as 1mM. The concentration of protein is also determined based on protein to lipid ratio (1uM for ratio of 1:1000). The residual OG in the mixture was removed by dialysis overnight in fusion buffer with SM2 bio-beads (Bio-Rad). The size of the vesicles was measured by dynamic light scattering (DLS), and the homogeneity of vesicles was also confirmed by checking size distribution using DLS (Supplementary Fig. 5).

### Single-molecule fluorescence assays

A quartz slide was cleaned by piranha solution (H_2_SO_4_:H_2_O_2_ = 2:1 in volume ratio) and 1M KOH (Chandradoss et al., 2014). Then, we coated the quartz slide with mPEG and biotin-PEG (LaySan Bio) in a 40:1 molar ratio, and assembled the slide with a coverslip as microfluidic chamber for total internal reflection (TIR) fluorescence microscopy. For all single molecule experiments, the quartz slides were washed with fusion buffer after every incubation to remove unbound reagents. To pulldown biotinylated vesicles, 0.1mg/ml Neutravidin (Thermo Fisher Scientific) was incubated on the quartz slide for 5 min and then proteo-liposomes were incubated for 5 min. For single-molecule FRET experiment characterizing the state of Sso1 opening, double labelled (Cy3, Cy5) Sso1 (Q120C/N215C) was embedded in the vesicles. For experiment to check Sec9 binding, Cy3-labeled Sec9 was incubated on the slide for the indicated times. To count fluorescent spots, TIR fluorescence images were recorded with an EM-CCD camera (iXON DU897D, Andor) under illumination with 532-nm laser (Spectra-Physics). Exposure time for the camera is 0.1s and this time is same for other single fluorescence imaging asaays.

For FRET experiment, fluorescence images were recorded for ~ 6min and data were collected. FRET efficiency E was calculated as follows.

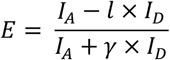

*I_A_* and *I_D_* are donor and acceptor intensities corrected with background signal. *l* is leakage and *γ* is gamma factor. Also, to ensure labeling of a single pair of Cy3 and Cy5, photobleaching was done for both Cy3 and Cy5 after the data was recorded. Using 532nm and 640nm laser with enhanced power, labeled Cy3 and Cy5 was bleached one by one. The alternating cycle of illumination with 532nm and 640nm laser is described in Supplementary Figure 1.

After we collected FRET values for all the recorded times, a histogram of FRET values was determined (the number of molecules analyzed was 43 for the experiment without exocyst and 33 with exocyst).

### Single vesicle docking and fusion assays

To detect the number of R-SNARE vesicles (DiD-labeled) pulled-down by binary Q-SNARE vesicles on the quartz slide via SNARE complex formation, we first mixed Sso1 Q-SNARE vesicles with Sec9 for 1 h at 30 °C to form the binary Sso1-Sec9 Q-SNARE complex. Then, R-SNARE vesicles were added to this solution, with or without exocyst. After incubation for certain amount of time (0, 15, 30, 60, 140 min), the mixture was incubated on the quartz slide. Finally, we recorded TIR fluorescence images with an EM-CCD camera under illumination with 640-nm laser.

To determine the extent of fusion of Q-SNARE vesicles and R-SNARE vesicles, PEG coated quartz slides were also prepared. The sample incubation process was the same as before except the incubation times and use of DiI-labelled Q-SNARE vesicles. The incubation times were 3, 7.5, 15,30,60,120 min. TIR fluorescence images were recorded with an EM-CCD camera under illumination with 532-nm laser and the calculations for FRET efficiency were the same as above.

### Experiments for single vesicle docking with exocyst

PEG coated quartz slides were also used for these experiments. After incubation of 0.1mg/ml Neutravidin, 0.01mg/ml biotinylated anti-eGFP antibody (Abcam) was incubated for 5 min. Then, the chamber was washed with BSA (Sigma-Aldrich) for reducing nonspecific interactions of vesicles to the quartz slide surface. eGFP-labeled exocyst complex was incubated for 10 min. Finally, DiD-labeled vesicles with individual SNAREs or SNARE complexes were incubated and there were no further washes. To check even distribution of biotinylated eGFP antibody for each set of experiments, fluorescence images of eGFP were recorded first under illumination with 488-nm laser. Then, we detected the fluorescence spot of DiD-labeled vesicles under illumination with 640nm laser. For analysis, we used two sample t-test. Five independent values were used for each set of experiment.

### Bulk vesicle fusion assays

The sample preparation process was the same as single vesicle fusion assays. Sso1 Q-SNARE vesicles (DiI-labeled) and Snc2 R-SNARE vesicles (DiD-labeled) were mixed to a final lipid concentration of 50μM. To measure the activity of vesicle lipid mixing, we measured acceptor fluorescence intensity by FRET using a plate reader (HIDEX Sense). The temperature was maintained at 30 °C during the experiment. We illuminated the mixture with a wavelength of 543nm for excitation of the donor (DiI) and detected the fluorescence signal at a wavelength of 680nm, for emission of the acceptor (DiD). Background intensity was measured separately and subtracted from the intensity of mixed vesicles.

### Content mixing assays

To make carboxy-fluorescein containing Snc2 R-SNARE vesicles, we added 100mM carboxy-fluorescein to protein-lipid mixture used for reconstitution of vesicles. This final mixture was dialyzed in fusion buffer with SM2 bio-beads and carboxy-fluorescein. After preparation of these vesicles, we prepared the quartz slides and pulldown Q-SNARE vesicles as described above. Then, we incubated carboxy-fluorescein contained Snc2 R-SNARE vesicles on the slide for 5 min. After washing away unbound vesicles and free carboxy-fluorescein, TIR fluorescence images were recorded with an EM-CCD camera under illumination with 488-nm laser (Supplementary Fig. 3).

### Dynamic light scattering (DLS)

To characterize sizes of the vesicles, a dynamic light scattering apparatus (Otsuka electronics ELSZ-1000) was used. 2mL of vesicle (0.2% (w/v)) in a glass-clear polystyrene cuvette (Ratiolab) was placed on the sample stage of the apparatus. Data were analyzed using the associated software (Otsuka electronics Photal).

## Acknowledgments

We thank M. Feyder and B. Reneker for technical assistance, and members of our labs for critical reading of the manuscript. Work in our laboratories is supported by National Creative Research Initiative Progeam NRF-2021R1A3B1071354 (T.-Y. Yoon), and National Institutes of Health grants GM068803 (M. Munson) and F32GM123704 (D. Lepore).

## References

Adamo, J.E., Rossi, G., and Brennwald, P. (1999). The Rho GTPase Rho3 has a direct role in exocytosis that is distinct from its role in actin polarity. Molecular biology of the cell 10, 4121–4133.

An, S.J., Rivera-Molina, F., Anneken, A., Xi, Z., McNellis, B., Polejaev, V.I., and Toomre, D. (2021). An active tethering mechanism controls the fate of vesicles. Nature communications 12, 1–14.

Antonin, W., Fasshauer, D., Becker, S., Jahn, R., and Schneider, T.R. (2002). Crystal structure of the endosomal SNARE complex reveals common structural principles of all SNAREs. Nature structural biology 9, 107–111.

Baker, R.W., Jeffrey, P.D., Zick, M., Phillips, B.P., Wickner, W.T., and Hughson, F.M. (2015). A direct role for the Sec1/Munc18-family protein Vps33 as a template for SNARE assembly. Science 349, 1111–1114.

Brennwald, P., Kearns, B., Champion, K., Keränen, S., Bankaitis, V., and Novick, P. (1994). Sec9 is a SNAP-25-like component of a yeast SNARE complex that may be the effector of Sec4 function in exocytosis. Cell 79, 245–258.

Bykhovskaia, M., Jagota, A., Gonzalez, A., Vasin, A., and Littleton, J.T. (2013). Interaction of the complexin accessory helix with the C-terminus of the SNARE complex: molecular-dynamics model of the fusion clamp. Biophysical journal 105, 679–690.

Carr, C.M., Grote, E., Munson, M., Hughson, F.M., and Novick, P.J. (1999). Sec1p binds to SNARE complexes and concentrates at sites of secretion. The Journal of cell biology 146, 333–344.

Chandradoss, S.D., Haagsma, A.C., Lee, Y.K., Hwang, J.-H., Nam, J.-M., and Joo, C. (2014). Surface passivation for single-molecule protein studies. JoVE (Journal of Visualized Experiments), e50549.

Chanturiya, A., Chernomordik, L.V., and Zimmerberg, J. (1997). Flickering fusion pores comparable with initial exocytotic pores occur in protein-free phospholipid bilayers. Proceedings of the National Academy of Sciences 94, 14423–14428.

D’Agostino, M., Risselada, H.J., Lürick, A., Ungermann, C., and Mayer, A. (2017). A tethering complex drives the terminal stage of SNARE-dependent membrane fusion. Nature 551, 634–638.

Diao, J., Ishitsuka, Y., Lee, H., Joo, C., Su, Z., Syed, S., Shin, Y.-K., Yoon, T.-Y., and Ha, T. (2012). A single vesicle-vesicle fusion assay for in vitro studies of SNAREs and accessory proteins. Nature protocols 7, 921–934.

Domanski, M., Molloy, K., Jiang, H., Chait, B.T., Rout, M.P., Jensen, T.H., and LaCava, J. (2012). Improved methodology for the affinity isolation of human protein complexes expressed at near endogenous levels. Biotechniques, 1.

Dubuke, M.L., Maniatis, S., Shaffer, S.A., and Munson, M. (2015). The exocyst subunit Sec6 interacts with assembled exocytic SNARE complexes. Journal of Biological Chemistry 290, 28245–28256.

Dubuke, M.L., and Munson, M. (2016). The secret life of tethers: the role of tethering factors in SNARE complex regulation. Frontiers in cell and developmental biology 4, 42.

Dulubova, I., Sugita, S., Hill, S., Hosaka, M., Fernandez, I., Südhof, T.C., and Rizo, J. (1999). A conformational switch in syntaxin during exocytosis: role of munc18. The EMBO journal 18, 4372–4382.

Fasshauer, D., Sutton, R.B., Brunger, A.T., and Jahn, R. (1998). Conserved structural features of the synaptic fusion complex: SNARE proteins reclassified as Q-and R-SNAREs. Proceedings of the National Academy of Sciences 95, 15781–15786.

Fukuda, R., McNew, J.A., Weber, T., Parlati, F., Engel, T., Nickel, W., Rothman, J.E., and Söllner, T.H. (2000). Functional architecture of an intracellular membrane t-SNARE. Nature 407, 198–202.

Ganesan, S.J., Feyder, M.J., Chemmama, I.E., Fang, F., Rout, M.P., Chait, B.T., Shi, Y., Munson, M., and Sali, A. (2020). Integrative structure and function of the yeast exocyst complex. Protein Science 29, 1486–1501.

Gao, Y., Zorman, S., Gundersen, G., Xi, Z., Ma, L., Sirinakis, G., Rothman, J.E., and Zhang, Y. (2012). Single reconstituted neuronal SNARE complexes zipper in three distinct stages. Science 337, 1340–1343.

Guo, W., Roth, D., Walch-Solimena, C., and Novick, P. (1999). The exocyst is an effector for Sec4p, targeting secretory vesicles to sites of exocytosis. The EMBO journal 18, 1071–1080.

Gurunathan, S., Chapman-Shimshoni, D., Trajkovic, S., and Gerst, J.E. (2000). Yeast exocytic v-SNAREs confer endocytosis. Molecular biology of the cell 11, 3629–3643.

Heider, M.R., Gu, M., Duffy, C.M., Mirza, A.M., Marcotte, L.L., Walls, A.C., Farrall, N., Hakhverdyan, Z., Field, M.C., and Rout, M.P. (2016). Subunit connectivity, assembly determinants and architecture of the yeast exocyst complex. Nature structural & molecular biology 23, 59–66.

Jun, Y., and Wickner, W. (2007). Assays of vacuole fusion resolve the stages of docking, lipid mixing, and content mixing. Proceedings of the National Academy of Sciences 104, 13010–13015.

Lehman, K., Rossi, G., Adamo, J.E., and Brennwald, P. (1999). Yeast homologues of tomosyn and lethal giant larvae function in exocytosis and are associated with the plasma membrane SNARE, Sec9. The Journal of cell biology 146, 125–140.

Lepore, D.M., Martínez-Núñez, L., and Munson, M. (2018). Exposing the elusive exocyst structure. Trends in biochemical sciences 43, 714–725.

Li, F., Tiwari, N., Rothman, J.E., and Pincet, F. (2016). Kinetic barriers to SNAREpin assembly in the regulation of membrane docking/priming and fusion. Proceedings of the National Academy of Sciences 113, 10536–10541.

Ma, C., Su, L., Seven, A.B., Xu, Y., and Rizo, J. (2013). Reconstitution of the vital functions of Munc18 and Munc13 in neurotransmitter release. Science 339, 421–425.

Ma, L., Rebane, A.A., Yang, G., Xi, Z., Kang, Y., Gao, Y., and Zhang, Y. (2015). Munc18-1-regulated stage-wise SNARE assembly underlying synaptic exocytosis. Elife 4, e09580.

MacDonald, C., Munson, M., and Bryant, N.J. (2010). Autoinhibition of SNARE complex assembly by a conformational switch represents a conserved feature of syntaxins (Portland Press Ltd.).

Mei, K., Li, Y., Wang, S., Shao, G., Wang, J., Ding, Y., Luo, G., Yue, P., Liu, J.-J., and Wang, X. (2018). Cryo-EM structure of the exocyst complex. Nature structural & molecular biology 25, 139–146.

Min, D., Kim, K., Hyeon, C., Cho, Y.H., Shin, Y.-K., and Yoon, T.-Y. (2013). Mechanical unzipping and rezipping of a single SNARE complex reveals hysteresis as a force-generating mechanism. Nature communications 4, 1–10.

Mollard, G.F.v., Nothwehr, S.F., and Stevens, T.H. (1997). The yeast v-SNARE Vti1p mediates two vesicle transport pathways through interactions with the t-SNAREs Sed5p and Pep12p. The Journal of cell biology 137, 1511–1524.

Morgera, F., Sallah, M.R., Dubuke, M.L., Gandhi, P., Brewer, D.N., Carr, C.M., and Munson, M. (2012). Regulation of exocytosis by the exocyst subunit Sec6 and the SM protein Sec1. Molecular biology of the cell 23, 337–346.

Munson, M. (2015). Synaptic-vesicle fusion: a need for speed. Nature structural & molecular biology 22, 509–511.

Munson, M., Chen, X., Cocina, A.E., Schultz, S.M., and Hughson, F.M. (2000). Interactions within the yeast t-SNARE Sso1p that control SNARE complex assembly. Nature structural biology 7, 894–902.

Munson, M., and Hughson, F.M. (2002). Conformational regulation of SNARE assembly and disassembly in vivo. Journal of Biological Chemistry 277, 9375–9381.

Nicholson, K.L., Munson, M., Miller, R.B., Filip, T.J., Fairman, R., and Hughson, F.M. (1998). Regulation of SNARE complex assembly by an N-terminal domain of the t-SNARE Sso1p. Nature structural biology 5, 793–802.

Ryu, J.-K., Min, D., Rah, S.-H., Kim, S.J., Park, Y., Kim, H., Hyeon, C., Kim, H.M., Jahn, R., and Yoon, T.-Y. (2015). Spring-loaded unraveling of a single SNARE complex by NSF in one round of ATP turnover. Science 347, 1485–1489.

Schwartz, M.L., Nickerson, D.P., Lobingier, B.T., Plemel, R.L., Duan, M., Angers, C.G., Zick, M., and Merz, A.J. (2017). Sec17 (α-SNAP) and an SM-tethering complex regulate the outcome of SNARE zippering in vitro and in vivo. Elife 6, e27396.

Scott, B.L., Van Komen, J.S., Irshad, H., Liu, S., Wilson, K.A., and McNew, J.A. (2004). Sec1p directly stimulates SNARE-mediated membrane fusion in vitro. The Journal of cell biology 167, 75–85.

Shen, D., Yuan, H., Hutagalung, A., Verma, A., Kümmel, D., Wu, X., Reinisch, K., McNew, J.A., and Novick, P. (2013). The synaptobrevin homologue Snc2p recruits the exocyst to secretory vesicles by binding to Sec6p. Journal of Cell Biology 202, 509–526.

Shon, M.J., Kim, H., and Yoon, T.-Y. (2018). Focused clamping of a single neuronal SNARE complex by complexin under high mechanical tension. Nature communications 9, 1–12.

Sivaram, M.V., Saporita, J.A., Furgason, M.L., Boettcher, A.J., and Munson, M. (2005). Dimerization of the exocyst protein Sec6p and its interaction with the t-SNARE Sec9p. Biochemistry 44, 6302–6311.

Song, H., Orr, A.S., Lee, M., Harner, M.E., and Wickner, W.T. (2020). HOPS recognizes each SNARE, assembling ternary trans-complexes for rapid fusion upon engagement with the 4th SNARE. Elife 9, e53559.

Strop, P., Kaiser, S.E., Vrljic, M., and Brunger, A.T. (2008). The structure of the yeast plasma membrane SNARE complex reveals destabilizing water-filled cavities. Journal of Biological Chemistry 283, 1113–1119.

Sutton, R.B., Fasshauer, D., Jahn, R., and Brunger, A.T. (1998). Crystal structure of a SNARE complex involved in synaptic exocytosis at 2.4 Å resolution. Nature 395, 347–353.

Togneri, J., Cheng, Y.-S., Munson, M., Hughson, F.M., and Carr, C.M. (2006). Specific SNARE complex binding mode of the Sec1/Munc-18 protein, Sec1p. Proceedings of the National Academy of Sciences 103, 17730–17735.

Tsui, M., and Banfield, D.K. (2000). Yeast Golgi SNARE interactions are promiscuous. Journal of Cell Science 113, 145–152.

Weber, T., Zemelman, B.V., McNew, J.A., Westermann, B., Gmachl, M., Parlati, F., Söllner, T.H., and Rothman, J.E. (1998). SNAREpins: minimal machinery for membrane fusion. Cell 92, 759–772.

Wiederkehr, A., De Craene, J.-O., Ferro-Novick, S., and Novick, P. (2004). Functional specialization within a vesicle tethering complex: bypass of a subset of exocyst deletion mutants by Sec1p or Sec4p. The Journal of cell biology 167, 875–887.

Yoon, T.-Y., and Munson, M. (2018). SNARE complex assembly and disassembly. Current Biology 28, R397–R401.

Yoon, T.-Y., Okumus, B., Zhang, F., Shin, Y.-K., and Ha, T. (2006). Multiple intermediates in SNARE-induced membrane fusion. Proceedings of the National Academy of Sciences 103, 19731–19736.

Yue, P., Zhang, Y., Mei, K., Wang, S., Lesigang, J., Zhu, Y., Dong, G., and Guo, W. (2017). Sec3 promotes the initial binary t-SNARE complex assembly and membrane fusion. Nature communications 8, 1–12.

Zhang, Y. (2017). Energetics, kinetics, and pathway of SNARE folding and assembly revealed by optical tweezers. Protein Science 26, 1252–1265.

