## Supplementary Figure for "Exocyst stimulates each step of exocytic SNARE complex assembly and vesicle fusion"

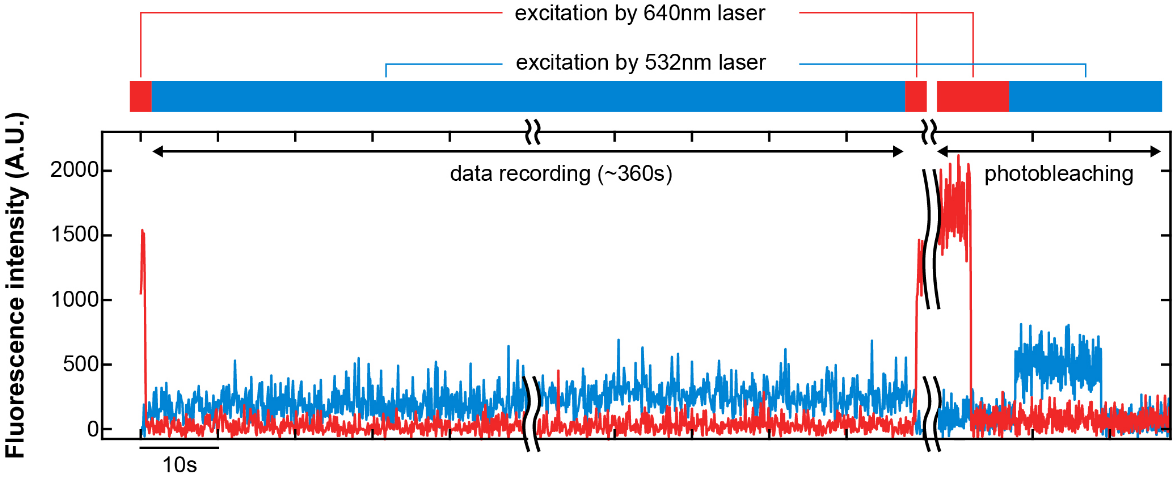


**Supplementary Fig 1. LASER illumination cycle for single-molecule FRET assay**

Fluorescence intensities of donor (green) and acceptor (orange) with respect to time. First, illumination by 640nm laser was performed for 0.5 s to check the existence of Cy5. Then, the data was recorded during illumination by 532nm laser. After that, the sample was illuminated by 640nm laser again to confirm that Cy5 was not photobleached. Lastly, for photobleaching labeled Cy3 and Cy5, the 640nm and 532nm lasers were turned on one at a time. The sudden decrease of intensity to background level by one step means photobleaching of a single dye.

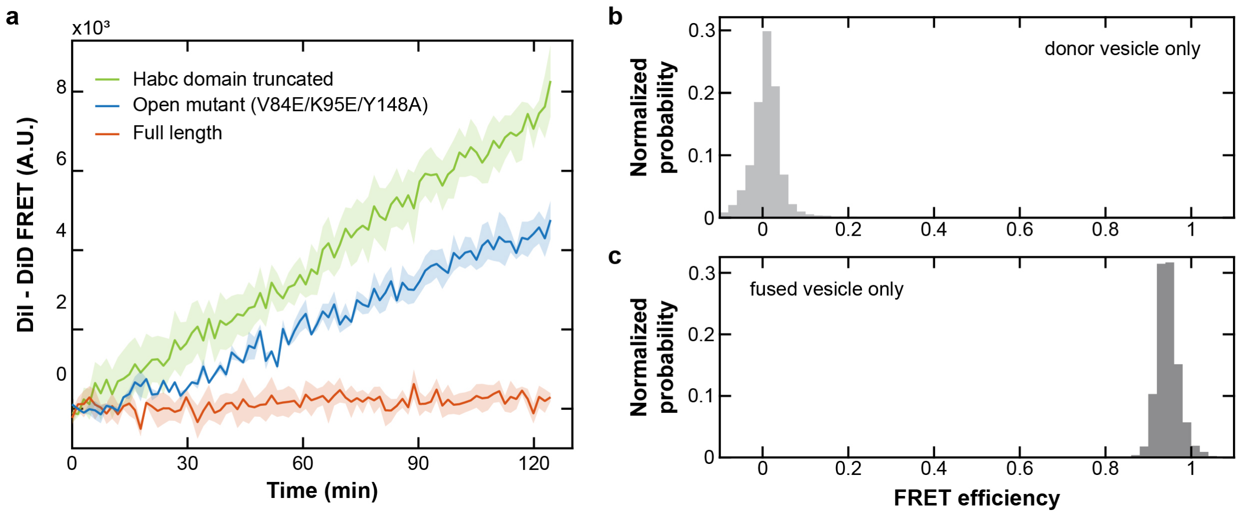


**Supplementary Fig 2. Lipid mixing assays with Sso1 open mutant and control data for identifying FRET values of lipid mixing**

**a**, DiI-DiD FRET signals generated by lipid mixing of Q-SNARE and R-SNARE vesicle. Three constructs of Sso1 was used: the Habc domain-truncated, the open mutant (V84E/K95E/Y148A) and full-length WT. **b**, FRET histogram of the DiI-labeled Q-SNARE vesicle alone. There was no lipid mixing and the FRET value was low (< 0.1). **c**, FRET histogram of fused vesicle alone. This vesicle had both DiI (2%) and DiD (2%) from adding both dyes in the vesicle reconstitution step, and demonstrates the vesicle signal upon completion of fusion process, where the FRET value is high (> 0.8).

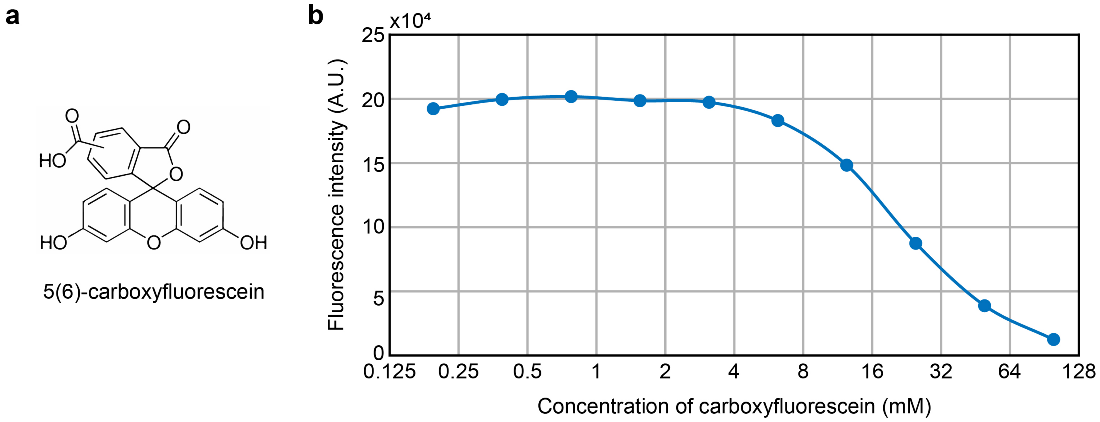


**Supplementary Fig 3. Fluorescence intensity of 5(6)-carboxyfluorescein**

**a,** Chemical structure of 5(6)-carboxyfluorescein. **b**, Fluorescence intensity of carboxyfluorescein becomes lower as the concentration increases, demonstrating the self-quenching property of carboxyfluorescein. The intensity was measured after diluting carboxyfluorescein in buffer, in the absence of vesicles.

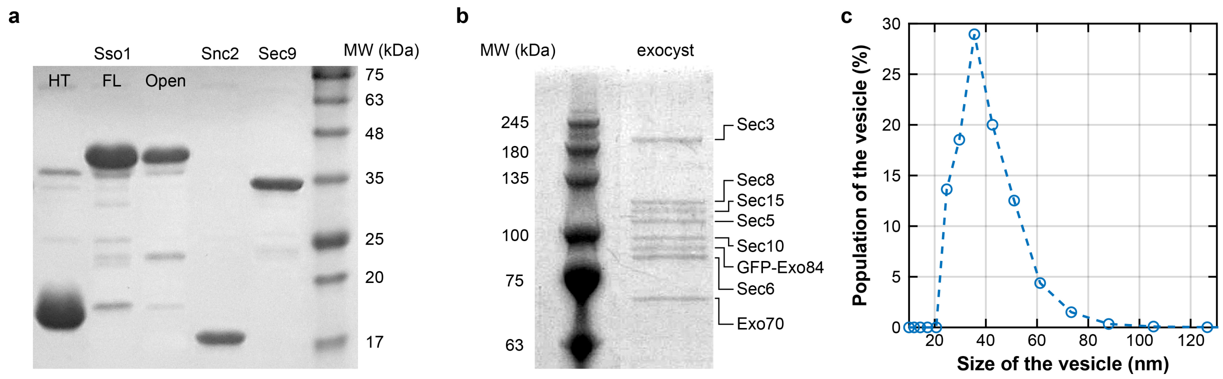


**Supplementary Fig 4. Purified yeast SNARE proteins and exocyst complex and characterization of vesicles**

**a**, Purified SNARE proteins analyzed by SDS-PAGE with Coomassie blue staining. 'HT' means Habc domain-truncated Sso1. 'FL' means full-length Sso1. 'Open' means Sso1 with mutations (V84E/K95E/Y148A). Molecular weight standards were in the rightmost lane. **b**, Purified exocyst complexes were analyzed by SDS-PAGE with Coomassie blue staining. Molecular weight standards were in the left lane. **c**, Size distribution of the vesicle measured by dynamic light scattering. The vesicle has same lipid composition that used for bulk vesicle fusion assay. Peak position is 35nm.
